# Feature selection by replicate reproducibility and non-redundancy

**DOI:** 10.1101/2023.07.04.547623

**Authors:** Tümay Capraz, Wolfgang Huber

## Abstract

**Motivation:** A fundamental step in many analyses of high-dimensional data is dimension reduction. Two basic approaches are introduction of new, synthetic coordinates, and selection of extant features. Advantages of the latter include interpretability, simplicity, transferability and modularity. A common criterion for unsupervised feature selection is variance or dynamic range. However, in practice it can occur that high-variance features are noisy, that important features have low variance, or that variances are simply not comparable across features because they are measured in unrelated numeric scales or physical units. Moreover, users may want to include measures of signal-to-noise ratio and non-redundancy into feature selection.

**Results:** Here, we introduce the RNR algorithm, which selects features based on (i) the reproducibility of their signal across replicates and (ii) their non-redundancy, measured by linear independence. It takes as input a typically large set of features measured on a collection of objects with two or more replicates per object. It returns an ordered list of features *i*_1_, *i*_2_, …, *i*_*k*_, where feature *i*_1_ is the one with the highest reproducibility across replicates, *i*_2_ that with the highest reproducibility across replicates after projecting out the dimension spanned by *i*_1_, and so on. Applications to microscopy based imaging of cells and proteomics experiments highlight benefits of the approach.

**Availability:** The RNR method is implemented in the R package *FeatSeekR* and is available via Bioconductor (Huber *et al*., 2015) under the GPL-3 open source license.

## 1 Introduction

Many biological datasets can be represented as a numeric matrix whose rows correspond to measured features and columns to objects of interest (e.g., cells, biological specimen). Here we consider settings where for each object we have two or more replicate measurements. Examples include RNA-Seq transcriptomics, mass spectrometry proteomics and microscopy-based cell morphology, where the features are levels of transcripts or proteins, or morphological descriptors of shape and texture of cells or cell compartments. The number of features can be in the thousands, but typically not all of them are informative (some are dominated by noise), and some are redundant of each other (they measure essentially the same underlying, relevant variable, in different ways). In this case, it can be desirable to reduce the dimensionality of the data.

Dimensionality reduction can be considered in supervised and unsupervised settings. Here we focus on the latter. There are two basic, not necessarily mutually exclusive, approaches: one is to introduce a smaller number of new variables that are linear or non-linear functions of the original variables; the other is feature selection. Examples for the first approach employ singular value decomposition (SVD), principal component analyis (PCA) (Jolliffe, 1986), and numerous versions of (non-linear) multi-dimensional scaling. As the new variables are smooth functions of the original features, random noise can cancel out. Sometimes they are meaningful “latent” variables. Here, however, we focus on feature selection, which can facilitate interpretation and integration of multiple datasets, and is attractively simple.

### 1.1 Related work

Unsupervised feature selection can be broadly categorized into embedded and filter methods. Embedded methods incorporate feature selection into the model fitting process and can be both supervised and unsupervised. An example for an unsupervised embedded method is sparse clustering where a penalization term is added to the clustering objective function (Witten and Tibshirani, 2010). Feature selection based on filtering uses properties of the data to prioritize features. Typically, features are ranked according to a statistic and the user defined top *n* features are selected. Different statistics of the data were previously used. These include mutual information and variance (Ferreira and Figueiredo, 2012; Guyon and Elisseeff, 2003), entropy (Varshavsky *et al*., 2006) or methods that minimize the reconstruction error of a projection of the original features induced by a selected subset of features (Wang *et al*., 2015). Here we introduce FeatSeekR, an unsupervised feature selection filter method, which uses replicate reproducibility as selection criterion. We were motivated for this work by Fischer *et al*. (2015), who devised a special case of our current method to use it on microscopy data, but only cursorily mentioned it in the supplement of their paper, without self-contained description, validation or software.

## 2 Approach

We posit that features carrying scientifically important information should be correlated between replicates. The algorithm iteratively selects features with the highest reproducibility across replicates, after projecting out those dimensions from the data that are linearly spanned by the previously selected features. Thus, each newly selected feature has a high degree of uniqueness.

## 3 Methods

The method pursues two aims. First, it selects features with high correlation between replicates, second, it aims to select features that are non-redundant between each other. We propose the following iterative, greedy procedure.

### 3.1 The FeatSeekR algorithm

Let **X** ∈ ℝ ^*pn*^ be a *p × n* data matrix with *n* observations each for *p* real-valued features. The columns of **X** represent repeated measurements on *k < n* biological conditions, or *k < n* different objects. The replication structure is encoded by the *n*-vector **r** ∈ {1, …, *k*}^*n*^, such that {*j* | *r*_*j*_ = *c*} are the indices of those columns in **X** that contain measurements for the *c*-th condition. For instance, if each condition was measured twice and replicates are next to each other in **X**, then **r** = (1, 1, 2, 2, 3, 3, …). We assume that most conditions have two or more replicates, but conditions with only one replicate are permitted. **X** may contain a small fraction of observations missing at random.

We label the iterations of the algorithm by the index *t* = 0, 1, 2, 3, … and denote by *S*_*t*_ the set of features selected up until iteration *t*. Thus, the elements of *S*_*t*_ are integers from 1 to *p*. Its complement 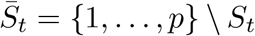 is the set of features not selected up until iteration *t*. The algorithm is greedy, so *S*_*t*_ ⊂ *S*_*t*+1_. The initial selection *S*_0_ is either the empty set ∅, or a set of features already pre-selected by the user based on criteria of their choice.

In iteration step *t*, we fit a linear model for each not previously selected feature 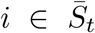 as a function of the selected features

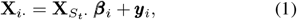

where **X**_*i·*_ is the *i*-th row of **X**, containing the observations of feature *i*, and 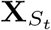 is the |*S*_*t*_| *× n* matrix obtained by subsetting from **X** the rows corresponding to *S*_*t*_. 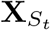 *·* contains the already selected features. 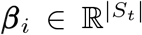 is the vector of coefficients for the regression of feature *i* on 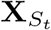 _*·*_, and ***y***_*i*_ ∈ ℝ^*n*^ the vector of residuals. We fit the free parameters on the right hand side of Eqn. (1) by linear regression, i.e., by minimizing the *L*_2_-norm of ***y***_*i*_.

We then use the residuals ŷ_*i*_ to represent the current (i.e., at step *t*) *non-redundant* information contributed by feature *i*. To measure replicate reproducibility of this non-redundant information, we use the *F* -statistic

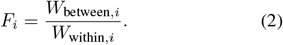

To compute *F*_*i*_, first define the overall mean 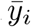 and the mean 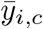 across replicates within condition *c* of 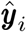:

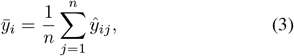

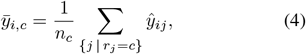

where *n*_*c*_ = |{*j* | *r*_*j*_ = *c*}| is the number of replicates for condition *c*, and we have hidden the dependence of these quantities on *t* in Equations (1)–(6) to unclutter the notation. Numerator and denominator of the *F* -statistic (2) are then:

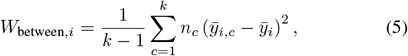

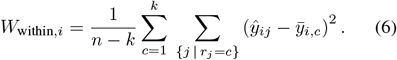

At the end of iteration step *t*, we select the feature *i*^*\**^ with highest *F*_*i*_ and proceed to the next iteration with *S*_*t*+1_ = *S*_*t*_∪{*i*^*^} until the user defined maximum number of selected features.

This procedure provides us with a list of features ranked by reproducibility and non-redundancy. A pseudocode representation is given in Algorithm 1.

### 3.2 Evaluation selected of feature subsets by fraction of explained variance

Optionally, to inform data-adaptive stopping in lieu of a predetermined value for the number of selected features, we can consider the fraction of variance of the dataset that is explained by the currently selected feature subset. We first model each feature **X**_*i·*_ of the original dataset **X** as a function of the selected features 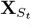 analogous to Equation (1). We then get the fraction of explained variance 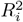 of each feature **X**_*i·*_ by calculating:

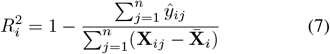

where 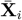 is the mean of feature *i*. We finally get the fraction of variance explained of the whole dataset by averaging *R*^2^ over all features.

#### Algorithm 1

FeatSeekR algorithm

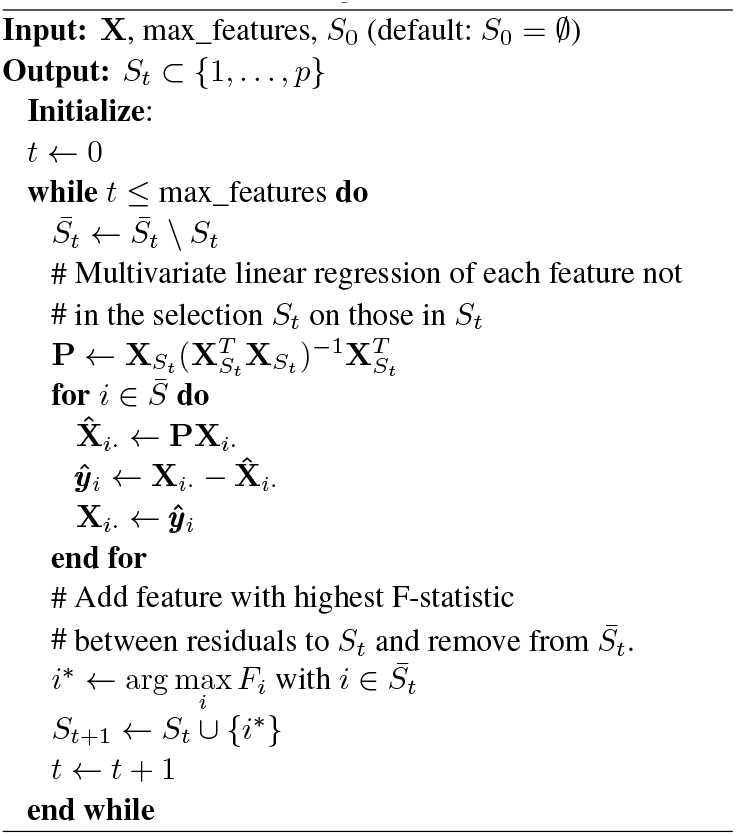

## 4 Results

### 4.1 Simulations

To demonstrate the algorithm we generated two synthetic datasets. The first dataset was characterized by a small number of underlying, “latent” variables that were noisily measured each by several observed features. In the second case we simulated data for a two-class clustering problem and compared our method to variance-based feature selection.

#### 4.1.1 Selecting non-redundant features

We generated an *l* × (*n*/3) matrix **M** by drawing each element *M*_*ij*_ independently from the standard normal distribution; *l* = 5 represents the number of groups and *n/*3 = 500 the number of objects. We applied the Gram-Schmidt process to orthonormalize the rows of **M**, resulting in an orthonormal matrix **Q**.

Next, for each group *i* (*i* ∈ 1 … *l*), we generated redundant features by scaling **Q**_*i·*_ by each of 10 random numbers *α*_*ij*_ ∼ *𝒩* (0, 1) (*j* ∈ 1 … 10) drawn independently from the standard normal distribution, i.e.: *X*_10(*i*−1)+*j,·*_ = *α*_*ij*_**Q**_*i·*_. This process yielded a 50 *×* 500 matrix we denote as **Q**^*t*^.

Finally, we created three replicates of **Q**^*t*^ by adding random numbers from the standard normal distribution element-wise to **Q**^*t*^, three times. The three replicates were concatenated, resulting in a final 50 *×* 1500 matrix **X**.

Figure 1 shows the correlation matrices of **X** and of the first five features selected by FeatSeekR. This result indicates that the algorithm is able to identify non-redundant features in this synthetic setting.

**Fig. 1.**
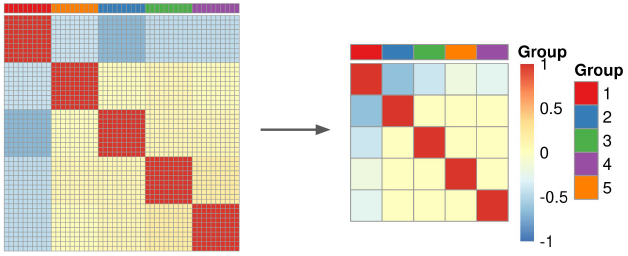
Left: correlation matrix of simulated data. Right: correlation matrix of first 5 selected features.

#### 4.1.2 Finding informative features in two-class data

We generated a *p* × *n* data matrix **X** ∈ ℝ^*p×n*^, where *n/*2 = 500 observations were divided into two classes and *p* = 50 features exhibit distinct signal-to-noise ratios. The mean values were assigned as follows: *μ*_1_, …, *μ*_*n/*2_ = 1 for observations in class 1, and *μ*_*n/*2+1_, …, *μ*_*n*_ = 2 for observations in class 2. For each feature *i*, we generated a vector **X**_*i·*_ ∼ *𝒩* (***μ***, *σ*_*i*_), where *σ*_*i*_ increased linearly as a function of *i* from 0.1 to 2. In this simulation setting, the two classes serve as replicates.

We ranked the features using two methods: FeatSeekR and based on their variance. We evaluated the feature selection via the performance of subsequent *k*-means clustering with *k* = 2 in recovering the two classes. For this, we calculated the adjusted Rand index (Morey and Agresti, 1984) between the clustering result and the known class labels.

Figure 2 shows the adjusted Rand index as a function of number of selected features. As might be expected, in each case the performance improves with increasing number of features, as that increasingly allows the noise to cancel itself out. However, selection by FeatSeekR achieves the same with a smaller number of selected features than the variance based selection. This result shows that feature selection based on total variance is not always an optimal criterion, as it conflates signal and noise, whereas FeatSeekR can disentangle these (see also Figure S1).

**Fig. 2.**
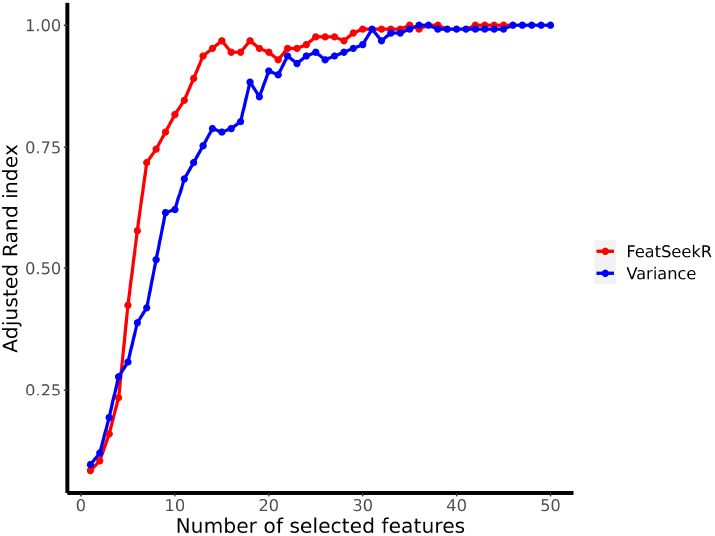
Performance of feature subsets selected by FeatSeekR and based on variance.

### 4.2 Applications to biological datasets

#### 4.2.1 Microscopy based image data from combinatorial knockout screens

Generic feature sets for microscopy based cytometry sets try to cover a wide range of information, ranging from general features such as intensity quantiles, object shapes (McQuin *et al*., 2018; Pau *et al*., 2010) or more abstract textural features introduced by (Haralick *et al*., 1973). As the produced features are designed to cover as much general information as possible, redundancy can be high. Additionally, not all features capture relevant information in every type of experiment. Consequently, some features are dominated by fluctuations that are irrelevant for the assay at hand, and not reproducible between repeated measurements. Here we used FeatSeekR to identify unique features with reproducible signal between measurements in two biological image datasets.

We used data from (Laufer *et al*., 2013), who performed combinatorial gene knock-downs in human cells using siRNA, followed by imaging, segmentation and feature extraction using the R package EBImage (Pau *et al*., 2010). A summary of datasets we used is shown in Table 1.

**Table 1.**
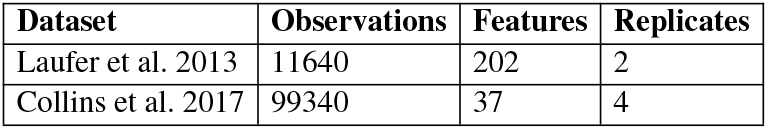
Summary of the used biological datasets.

Analogous to our idealized example in Figure 1, the extracted features formed groups of high correlation within and lower correlation between (Figure S2). The grouping was partially interpretable as some groups broadly corresponded to different colour channels (fluorescent labels) or cellular compartments. This supports the idea that the effective dimension of the data matrix is substantially lower than the number of features *p* = 202 and that feature selection is a plausible approach to these data. We used FeatSeekR to select a set of features that explained more than 70% of the variance of the original dataset. The selection comprised of 5 features, a substantial reduction (Figure 3A). The overall low correlation between the selected features confirmed their low redundancy. We note that selected feature sets are dependent on the starting set of features. For instance, the features *Cell actin majoraxis* and *Cell actin eccentricity* are highly correlated, and when calling FeatSeekR with preselection of *Cell actin eccentricity, Cell actin majoraxis* was not selected. This illustrates that multiple selections are equally admissible and can be influenced by a user-defined preselected set of features.

**Fig. 3.**
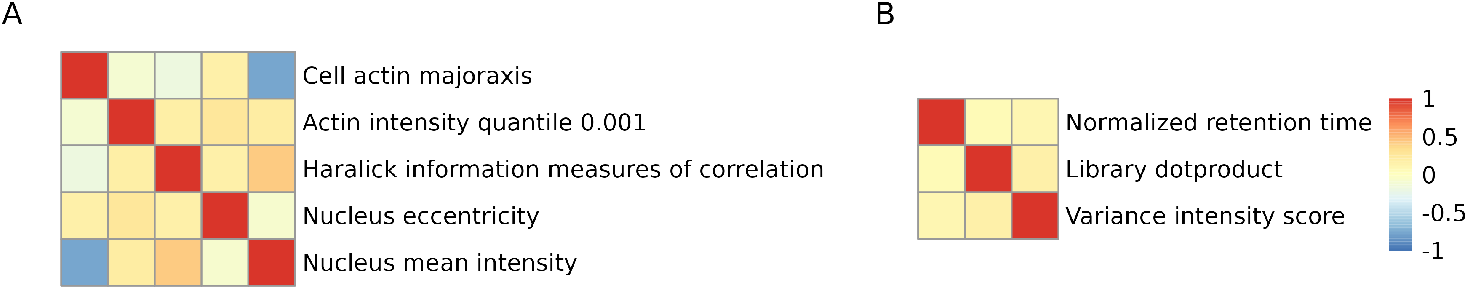
Correlation matrix of selected features of A (Laufer et al., 2013) and B (Collins et al., 2017) that explain at least 70% of the variance of the original dataset. Features are colored according to their feature clusters.

#### 4.2.2 Mass spectrometry-based proteomics data

Next we applied FeatSeekR to spectral features of a proteomics dataset from (Collins *et al*., 2017), where the authors investigated the reproducibility of a mass spectrometry based proteomics measurement across multiple international sites. In this type of experiment, proteins are usually first digested to peptides, separated via LC-MS and their mass spectra are subsequently recorded. To identify individual peptides, mass spectra are either matched to a database or to a library of spectra of known peptides (Aebersold and Mann, 2016). Beforehand, features such as retention time, intensities and mass accuracies are extracted. The matching to the reference is then done based on these extracted features (Röst *et al*., 2014). We used measurements of 4 sites as replicates, leading to 99340 peptide assays (observations), 37 features and 4 replicates (see Table 1).

We observed that not all of these automatically extracted spectral features are equally reproducible and informative across sites. Furthermore, the dataset consists of several correlated redundant feature clusters. For example peak features related to retention time, distance to the reference library or p-value related features form very distinct clusters (Figure S3). We used FeatSeekR to select the most reproducible features that explained at least 70% of the total variance. Figure 3B shows that we identified the most reproducible features of the redundant and correlated feature clusters. The selected features cover both peptide retention time, as well as information related to their mass spectra.

## 5 Discussion

We present a framework for feature selection which selects features based on their reproducibility between replicates while keeping redundancy low. In contrast to existing filtering based feature selection methods, we make use of replicated measurements and are able to effectively separate biological signal from noise. Additionally, FeatSeekR is capable of performing feature selection on ragged data, where not all conditions or observed objects have the same number of replicate observations. We show on synthetic data that FeatSeekR is able to find exactly one feature per underlying latent factor. We highlighted its utility as a preprocessing step for clustering, by selecting more informative features and removing more noisy ones. Furthermore, we show the application of our method to biological data, derived from microscopy based imaging of cells and proteomics experiments. Our algorithm finds feature sets of biological datasets that achieve a good trade-off between captured information and redundancy.

In practice, feature selection can serve different purposes, such as reduction of storage space and computation time, or better performance of downstream machine learning methods. If FeatSeekR is used to improve performance in a machine learning context, feature selection should be incorporated in the cross validation procedure (Ambroise and McLachlan, 2002).

To guide the selection process, we provide diagnostic tools to analyze and visualize information content in biological datasets, within the FeatSeekR package.

The objective that motivates feature selection with FeatSeekR does not lead to a unique optimal selection. Conceptually, it is compatible with multiple selections that are, for practical purposes, equally admissible. Thus, even if the implementation by FeatSeekR returns a single selection, this should be viewed as a representative proposal, not as a unique solution. FeatSeekR uses a greedy algorithm and is not based on a global optimality criterion. Formulating such a global optimality criterion and associated algorithms remains a direction for future research.

## Acknowledgements

This work was funded by the European Research Council (ERC Synergy Project DECODE under grant agreement no. 810296).

## 6 Supplementary Figures

**Fig. S1.**
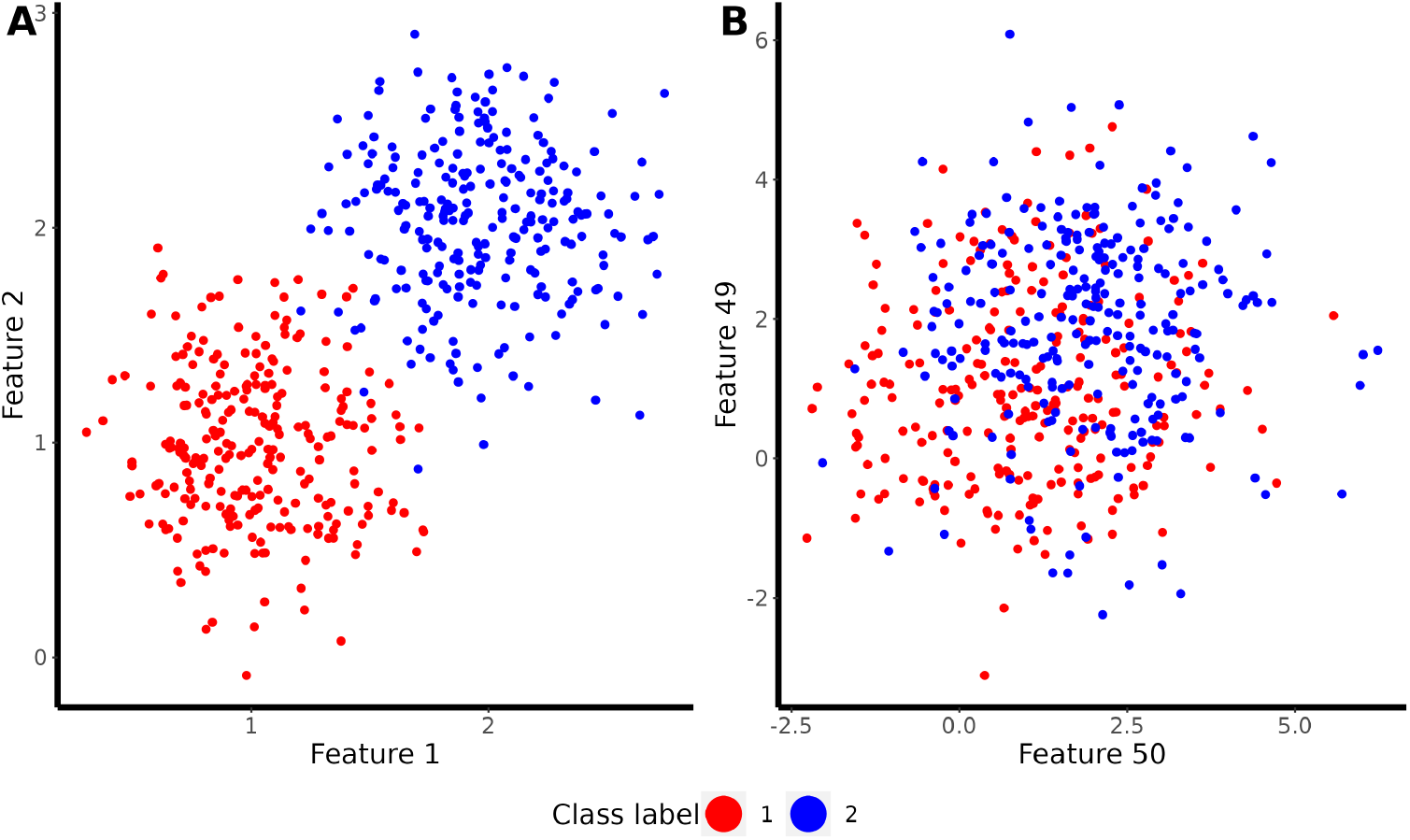
Scatter plots of simulated 2 class data. Features in (A) have lower variance than in (B) and show better class separation.

**Fig. S2.**
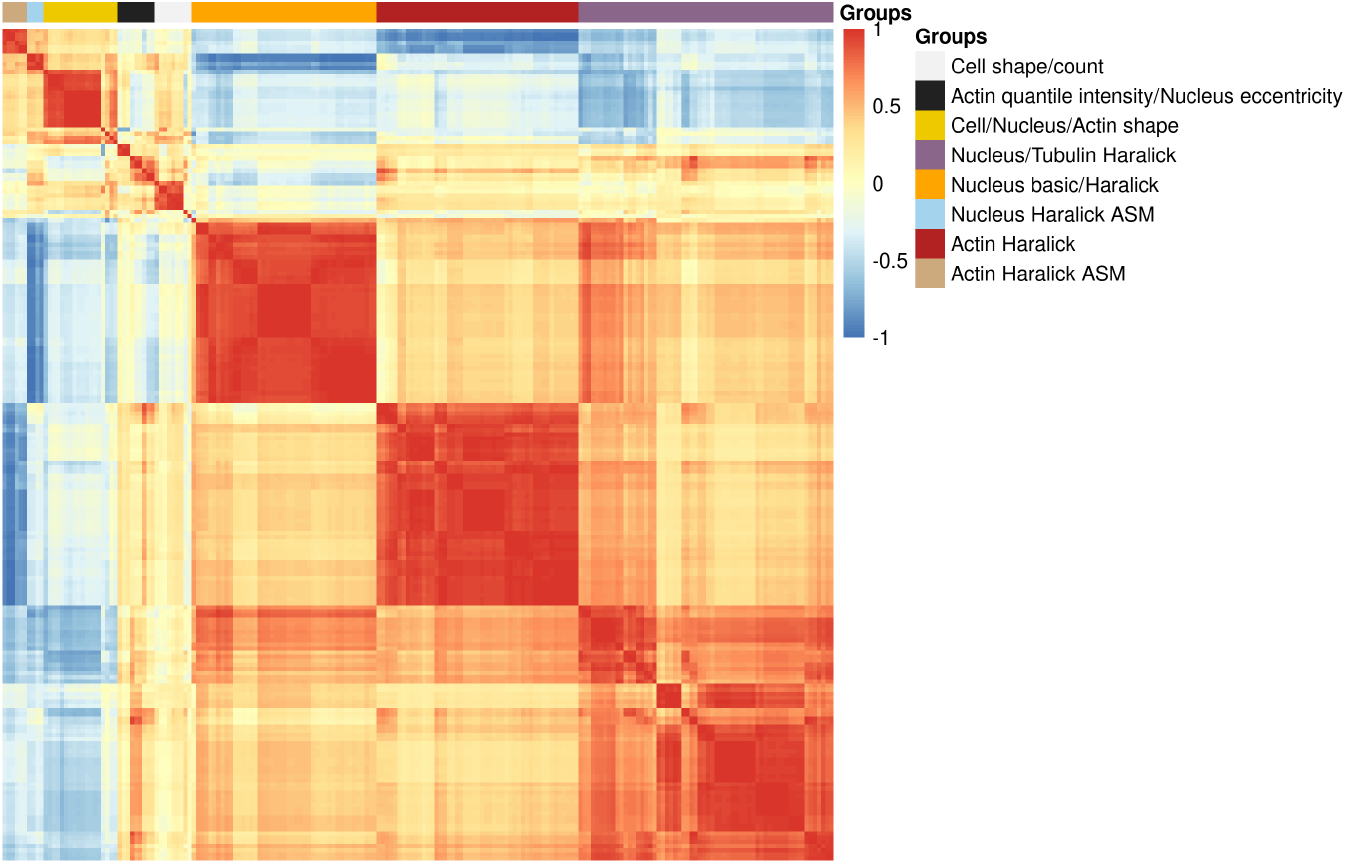
Correlation matrix of the features from (Laufer et al., 2013)

**Fig. S3.**
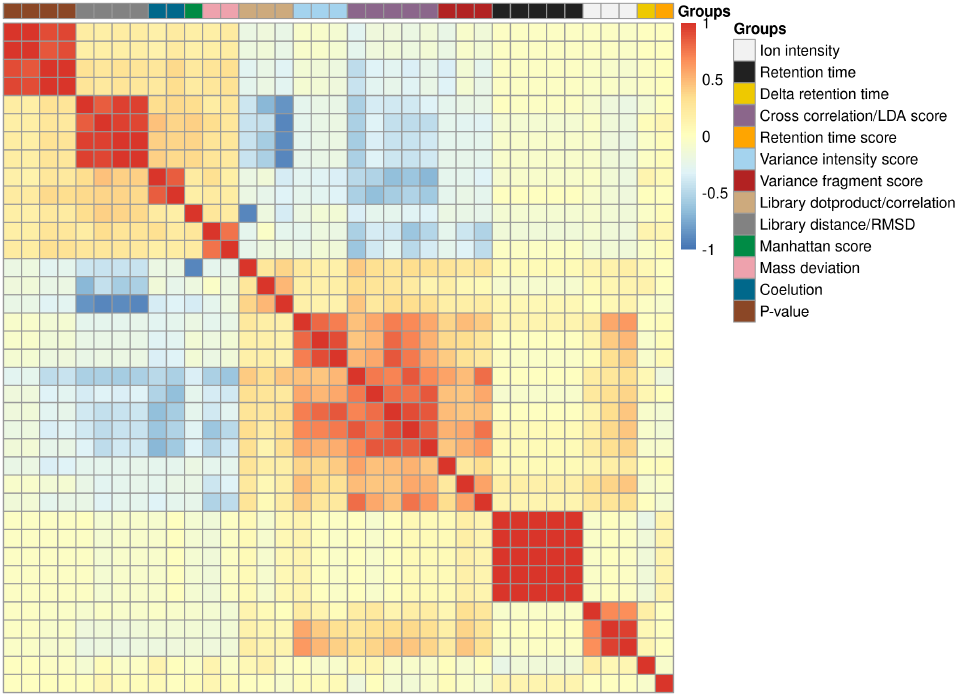
Correlation matrix of the features from (Collins et al., 2017)

